# Effects of food supplementation and helminth removal on space use and spatial overlap in wild rodent populations

**DOI:** 10.1101/2022.12.22.521674

**Authors:** Janine Mistrick, Jasmine S.M. Veitch, Shannon M. Kitchen, Samuel Clague, Brent C. Newman, Richard J. Hall, Sarah A. Budischak, Kristian M. Forbes, Meggan E. Craft

**Affiliations:** Department of Ecology, Evolution, and Behavior, University of Minnesota, St. Paul, MN, USA; W.M. Keck Science Department, Claremont McKenna, Pitzer, and Scripps Colleges, Claremont, CA, USA; Department of Biological Sciences, University of Arkansas, Fayetteville, AR, USA; Odum School of Ecology, University of Georgia, Athens, GA, USA; Department of Infectious Diseases, College of Veterinary Medicine, University of Georgia, Athens, GA, USA; Center for the Ecology of Infectious Diseases, University of Georgia, Athens, GA, USA

**Keywords:** anthelmintic treatment, bank vole, *Clethrionomys glareolus*, food supplementation, network analysis, space use, spatial overlap

## Abstract

1. Animal space use and spatial overlap can have important consequences for population-level processes such as social interactions and pathogen transmission. Identifying how environmental variability and inter-individual variation affect spatial patterns and in turn influence interactions in animal populations is a priority for the study of animal behavior and disease ecology. Environmental food availability and macroparasite infection are common drivers of variation, but there are few experimental studies investigating how they affect spatial patterns of wildlife.

2. Bank voles (*Clethrionomys glareolus*) are a tractable study system to investigate spatial patterns of wildlife and are amenable to experimental manipulations. We conducted a replicated, factorial field experiment in which we provided supplementary food and removed helminths in vole populations in natural forest habitat and monitored vole space use and spatial overlap using capture-mark-recapture methods.

3. Using network analysis, we quantified vole space use and spatial overlap. We compared the effects of food supplementation and helminth removal and investigated the impact of season, sex, and reproductive status on space use and spatial overlap.

4. We found that food supplementation decreased vole space use while helminth removal increased space use. Space use also varied by sex, reproductive status, and season. Spatial overlap was similar between treatments despite up to three-fold differences in population size.

5. By quantifying the spatial effects of food availability and macroparasite infection on wildlife populations, we demonstrate the potential for space use and population density to trade off and maintain consistent spatial overlap in wildlife populations. This has important implications for spatial processes in wildlife including pathogen transmission.

## 1 INTRODUCTION

How animals use space in their environment and interact with conspecifics is fundamental to understanding proximity-and contact-driven processes such as social interactions (Kusch & Lane, 2021) and pathogen transmission (Silk et al., 2018). Interactions with conspecifics are frequently heterogeneous, reflecting individual variation due to sex and reproductive status and associated social behavior or, alternatively, can be driven by environmental variability in habitat quality or population density (Craft, 2015; Tompkins et al., 2011). Since animal space use and conspecific interactions are crucial determinants of pathogen exposure, identifying how these are shaped by individual-level variation and environmental variability has important implications for mitigating effects of novel and zoonotic pathogens on wildlife, domestic animals, and humans (Silk et al., 2017). However, studies that disentangle the relative contributions of environmental and individual variation to interactions between animals are rare.

Food availability is a common source of environmental variability that can alter animal space use and spatial overlap and influence pathogen transmission (Becker et al., 2015). At local scales, low food availability may increase space use through increased foraging, while areas of high food availability may decrease space use through increased site fidelity; heterogeneity in food availability across the landscape may promote movement through increased attraction and dispersal between patches (Becker et al., 2018). Point food sources (e.g., bird feeders, landfills, crop fields) can increase spatial overlap of wildlife through aggregation and facilitate both direct interactions between infected and susceptible animals (Altizer et al., 2018; Forbes et al., 2015) and indirect interactions through environmentally transmitted pathogens accumulated in the environment (Cross et al., 2007).

Individual variation among animals can also affect space use and spatial overlap with potential consequences for pathogen transmission (VanderWaal & Ezenwa, 2016). For example, in rodents, testosterone-mediated behavior in mature males can increase their contacts with conspecifics and transmission potential (Grear et al., 2009) and is hypothesized to be a key driver of commonly observed male bias in infection prevalence (Krasnov et al., 2012). Individual variation may also arise through differing burdens of macroparasites such as helminths, with potentially opposing effects on space use and spatial overlap. Helminth infection could increase feeding behavior, requiring infected hosts to expand their foraging area (Brown et al., 1994). Conversely, helminths may cause sickness-induced lethargy in infected animals and decrease their activity, decreasing their space use and interactions with conspecifics (Ghai et al., 2015). Further, since space use and spatial overlap can influence macroparasite exposure, positive feedbacks between macroparasite exposure and space use could exacerbate or regulate within-population variation in macroparasite burden and movement (Hawley et al., 2021).

Much of our knowledge of how food availability and macroparasite infection impact transmission in wildlife comes from studies of laboratory animals under controlled conditions (Croft et al., 2011) or observational studies in wildlife (Jolles et al., 2008) and therefore often lack explicit investigation linking spatial behavior to transmission potential. In natural settings, food availability and macroparasite infection frequently co-vary and could have balancing or synergistic effects on animal space use and spatial overlap. For example, abundant food resources could alleviate increased foraging requirements caused by helminth infection, resulting in minimal effects of macroparasite burden on space use. Ultimately, rigorous field experiments in wildlife are necessary to test the independent and synergistic effects of environmental and individual variation on animal spatial patterns and implications for pathogen transmission (e.g., Sweeny et al., 2021). Such research is pertinent and timely because land-use change is altering patterns of food availability and macroparasite burdens that can exacerbate pathogen emergence and transmission at the human-wildlife interface (Plowright et al., 2021).

Our goal was to determine the effects of food supplementation and macroparasite removal on space use and spatial overlap in wildlife populations. We used bank voles (*Clethrionomys glareolus*, previously *Myodes glareolus*) as our study species as they are amenable to experimental manipulation, are a common host for macroparasite infection (e.g., helminths like *Heligmosomoides glareoli* can have a population-level prevalence of up to 80%; Haukisalmi & Henttonen, 2000), and have well-documented natural history, particularly with respect to spatial behavior (Tamarin et al., 1990). Reproductive males hold territories that are approximately double those of females and that overlap with territories of males and females (reproductive males average: 1800 m^2^; females: 900 m^2^; Bujalska & Grüm, 1989). Reproductive females are territorial toward other reproductive females but may overlap more in the non-breeding season (Bujalska, 1990). In both sexes, space use of non-reproductive voles is less than that of reproductive voles (non-reproductive females, males average: 700 and 800 m^2^, respectively; Bujalska & Grüm, 1989). Both food availability and population density are thought to regulate vole territory size (Bondrup-Nielsen & Karlsson, 1985).

We experimentally manipulated wild bank vole populations via food supplementation and macroparasite (helminth) removal and monitored individual-and population-level responses. Specifically, we 1) quantified space use by sex and reproductive status in the breeding and non-breeding seasons, and 2) quantified spatial overlap between voles each month. Space use and spatial overlap were compared between treatments to test the effects of food supplementation and helminth removal on vole spatial patterns. As abundant food often increases small mammal population density through immigration (Prevedello et al., 2017), we hypothesized that food supplementation would decrease vole space use and increase spatial overlap. We hypothesized that helminth infection would increase host movement (due to increased feeding behavior), and thus helminth removal would decrease space use and spatial overlap.

## 2 MATERIALS AND METHODS

### 2.1 Study site and experimental design

We conducted a two-factor field experiment in boreal forests of southern Finland (61.0775°N, 25.0110°E) where bank voles are the dominant rodent species. These forests receive snowfall from approximately November-April, and absence of snow cover makes monitoring and experimental manipulation of voles most feasible from May-October. Twelve study sites were established in old-growth spruce forest patches. All sites were at least two kilometers apart to prevent dispersal between replicate populations (maximum dispersal distance estimated to be 500 m in heterogeneous habitat, Gliwicz & Ims, 2000). At each site, a standardized trapping grid (100 m x 100 m, 1 hectare) was established to monitor the vole population, the grid consisting of 61 uniquely identified traps spaced 10 meters between grid rows and columns (Fig. 1).

**Figure 1.**
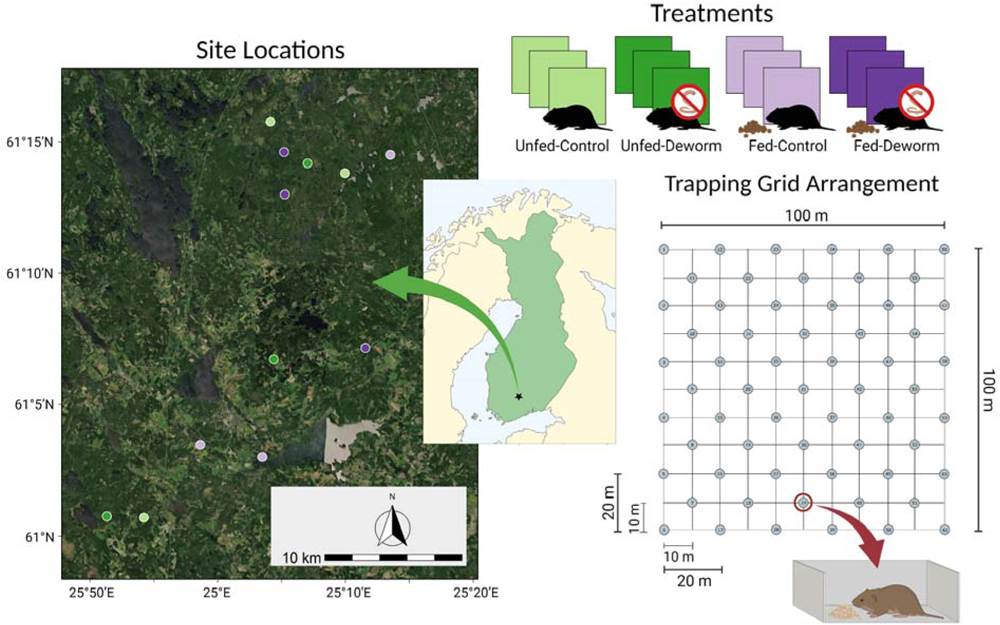
Field study design and experimental set-up. (“Site Locations”) Twelve field sites were established in southern Finland. (“Treatments”) Sites were assigned one of four treatment pairings: unfed-control, unfed-deworm, fed-control, and fed-deworm and each treatment was replicated at three sites each. (“Trapping Grid Arrangement”) The trapping grid included 61 traps in 11 rows and 11 columns. Rows and columns were spaced 10 meters apart and traps (dots) were placed in an offset arrangement with 20 meters to the next trap in the same row and column and 14.14 meters to the next trap on the diagonal.

Sites were randomly assigned to one of four treatment pairings (“treatments”): no manipulation (“unfed-control”; “U-C”); helminth removal only (“unfed-deworm”; “U-D”); food supplementation only (“fed-control”, “F-C”); and both food supplementation and helminth removal (“fed-deworm”; “F-D”). Each treatment was replicated at three sites (Fig. 1). Sites assigned to food supplementation received a feed mix of mouse chow pellets and sunflower seeds evenly distributed on the trapping grid every two weeks from May through November (if snowfall had not yet accumulated). The supplemental feed mix was dosed at 7.54 kcal/m^2^ (following Sweeny et al., 2021) consisting of 7.875 kg of mouse chow pellets (Altromin 1324 10 mm pellets maintenance diet for rats and mice [3227 kcal/kg]; Altromin Spezialfutter GmbH & Co. KG, Lage, Germany) and 7.875 kg sunflower seeds (as fed, 6350 kcal/kg). Voles at deworm sites were given an oral dose of 10 mg/kg Ivermectin and 100 mg/kg Pyrantel on their first capture in a trapping occasion; this combination of medication is effective at treating larval and adult helminth infections (Clerc et al., 2019). Voles at control sites received a matching weight-based control dose (17.5% sucrose solution) on their first capture in a trapping occasion.

### 2.2 Data collection

Vole populations were longitudinally monitored via capture-mark-recapture methods. Every four weeks from May-October 2021, Ugglan multi-capture live traps (Grahnab, Sweden) were baited with whole oats and set for 48 hours (a “trapping occasion”). Traps were checked each morning and evening at approximately 0600 and 1600 (four checks per trapping occasion) and captured animals were processed and then released at the capture location. Six trapping occasions were conducted at each site during the study period (total trap events: 17,568). Upon first capture, all voles were injected with a Passive Integrated Transponder (“PIT tag”; “Skinny” PIT Tag, Oregon RFID, USA) for unique identification. The trap number, PIT tag number, sex, and reproductive status were recorded and a fecal sample was collected (to quantify helminth presence and infection intensity via fecal egg counts) for each vole on their first capture each trapping occasion. If a vole was recaptured in a trapping occasion, only the trap number and PIT tag number were recorded, no samples were collected, and the vole was released at the capture location.

Each capture, voles were categorized as reproductive (perforate vagina, lactating, or pregnant females; males with scrotal testes) or non-reproductive. Reproductive status per vole was summarized within the summer breeding season (June-August) and autumn non-breeding season (September-October in southern Finland; Kaikusalo, 1972) to account for seasonal differences in vole behavior. If a vole was ever recorded as reproductive during a trapping occasion within the season, it was recorded as reproductive in that season. In autumn, voles that were reproductive in summer were also considered reproductive in autumn, even if external traits were not observed in September or October. Voles were further classified into four functional groups: reproductive males, reproductive females, non-reproductive males, and non-reproductive females. Functional groups are population subgroups that are relatively uniform in their behavior, physiology, and immunology. Conducting epidemiological analysis at this level pools individuals into biologically meaningful groups and enables more realistic conclusions to be drawn (Henttonen, 2022).

All trapping, handling, and sampling of wild bank voles was conducted under approval of the University of Arkansas Institutional Animal Care and Use Committee (IACUC #19105) and the Finnish Animal Ethics Board (ESAVI-17810-2019). Access to forest sites was provided by private landowners and by Metsähallitus Metsätalous Oy (MH 6302/2019).

### 2.3 Data analysis

The efficacy of the deworm treatment was assessed using helminth infection status and intensity in infected animals (eggs per gram of feces [EPG]), quantified using a salt flotation method (Pritchard & Kruse, 1982). Helminth infection status was modeled for all sampled animals using a generalized linear mixed-effects model (binomial family, logit link; “lme4” R package, Bates et al., 2015). Helminth intensity (natural log-transformed EPG) was modeled for infected animals using a linear mixed-effects model (lmerTest R package, Kuznetsova et al., 2017). Both models were parameterized with fixed effects of: treatment group (deworm/control), treatment stage (pre-treatment [the first capture of a vole] / post-treatment [all subsequent captures]), and the interaction of treatment group and stage. Vole ID was included as a random effect. Model interactions were visualized using the ‘visreg’ R package (Breheny & Burchett, 2017).

To understand general patterns of vole space use and how they were influenced by food supplementation and helminth removal, the mean space use of voles in each functional group was characterized in both the breeding and non-breeding seasons for each treatment. Within a single site, overlapping space use between pairs of voles (“pairwise space-use overlap”) was used to approximate opportunities for direct and indirect interactions (Robert et al., 2012). The total pairwise space-use overlap of a focal vole with all its neighbors (“individual spatial overlap”) was quantified to investigate how opportunities for interactions varied among individuals. When interactions like spatial overlap are heterogeneous among individuals, network analysis is a useful method to visualize their frequency and distribution within a population (Croft et al., 2008; Krause et al., 2007). We constructed spatial overlap networks to visualize all the pairwise space-use overlap between voles at each site to investigate how the experimental manipulations affected opportunities for direct and indirect interactions at the population level.

Vole capture numbers in May were very low (zero to five animals per site), prohibiting network construction for all sites, therefore May capture data were excluded from downstream analysis. Additionally, since space use was estimated by functional group (the combination of sex and reproductive status), only animals with both sex and reproductive status data recorded were included in the analysis.

Data analysis was conducted in program R version 4.1.2 (R Core Team, 2021).

#### 2.3.1. Factors influencing seasonal space use

Seasonal vole space use was approximated within a season by aggregating capture location data among individuals by population subgroups to characterize the average space use of an individual in each group (Wanelik & Farine, 2022). This approach allows space use and spatial overlap to be estimated for individuals, even if they are caught only once or only in one trap and helps minimize uncertainty in space-use estimates created by the relatively short observation periods and limited observations per individual. As such, these methods offer a robust way to detect biological effects of shared space use when using sparse capture-mark-recapture data where repeat observations per individual are limited or heterogeneous (Wanelik & Farine, 2022).

Using this approach, we aggregated vole capture locations from each trapping occasion June-October (hereafter “month”) into a summer breeding season (June-August) and an autumn non-breeding season (September-October). We defined a vole’s seasonal “centroid” as the weighted average of all locations where it was trapped that season, allowing multiple captures in the same trap to more strongly influence the centroid than a single capture in a trap. For each vole, the distance from this centroid to every trap in the trapping grid (n=61) was calculated. Then, distances for all voles (across the entire study) were pooled by season.

We used generalized linear models (GLMs) to characterize the probability of capturing a vole in a trap a given distance from its seasonal centroid and investigate factors influencing space use (i.e., sex, reproductive status, treatment), separately in summer and autumn. In each model, the response variable was whether a vole was caught in a given trap (where 1 indicated it was caught, 0 indicated it was not) and the explanatory variables were: the natural log distance of the trap from the vole’s seasonal centroid, vole sex, reproductive status (reproductive/non-reproductive), site-level food treatment (unfed/fed), helminth treatment (control/deworm), and interaction effects between distance and each of: sex, reproductive status, food treatment, and helminth treatment, and a three-way interaction between distance, food treatment, and helminth treatment. The interaction terms were examined to identify parameters that significantly affected the relationship between distance from the seasonal centroid and capture probability, indicating differences in space use. Model interactions were visualized using the ‘visreg’ R package.

#### 2.3.2. Seasonal space use

We then used the significant predictors of capture probability as identified by the seasonal space use GLMs to divide voles into population subgroups and characterize space use separately for each group. The distances from a vole’s seasonal centroid to every trap in the trapping grid were pooled across voles, grouped by the combination of season*sex*reproductive status*food treatment*helminth treatment (n groups = 32). For each group, we fitted a negative sigmoidal curve (Equation 1; fitted using a Bernoulli GLM, where 1 indicated an animal was caught in a given trap, 0 indicated it was not; as developed by Wanelik & Farine, 2022) to characterize the average space use of a vole in that group:

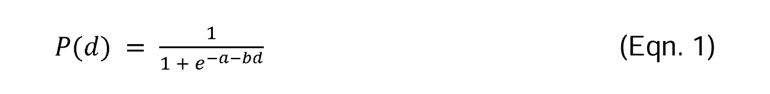

where space use was defined as the declining probability P of capturing a vole an increasing distance (d) from its seasonal centroid as determined by: a (describing the size of the space use radius), b (describing how steeply the capture probability declines as distance increases), and d (the logarithmic distance from the centroid). Thus, space use was separately characterized in the summer and autumn for each treatment for voles of each functional group.

#### 2.3.3. Pairwise space-use overlap and spatial overlap networks

Space-use overlap was estimated between all pairs of voles captured at a site in a given month. These pairwise interactions were then used to create spatial overlap networks to visualize all the interactions in the observed population at each site, in each month. First, the monthly centroid for every vole was calculated as the weighted average of all trap locations where it was captured that month (1-4 captures/month possible). The appropriate seasonal space use for a given vole’s sex, reproductive status, the site-level food and helminth treatments, and the season (as calculated above using Eqn. 1) was centered on this monthly centroid.

Overlap between all pairs of voles observed in a month at a site was estimated based on the overlap in the two voles’ space use (as developed by Wanelik & Farine, 2022). Specifically, we calculated the amount of pairwise space-use overlap by first predicting the (independent) probabilities of capturing vole 1 and vole 2 (using Eqn. 1) at each grid point (x, y) in a grid of coordinates X and Y overlapping both of their space use. The amount of pairwise space-use overlap between the two voles (£_1,2_) was estimated by taking the lower of the two voles’ capture probabilities at each grid point (x, y) and summing them across all grid points and similarly summing the higher of the two capture probabilities across all grid points. The summed minimums were then divided by the summed maximums to get a ratio indicating how similar the two voles’ capture probabilities were across space (Equation 2, developed by Wanelik & Farine, 2022):

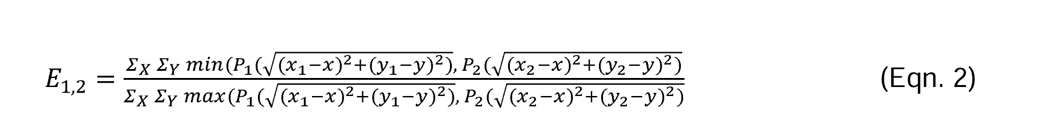

The more similar the space use of the two voles, the ratio (£_1,2_) approaches 1 (the two voles have similar capture probabilities at each grid point, indicating more complete overlap), whereas the more different their space use, the ratio approaches 0 (where one vole has high capture probability, the other has low capture probability, indicating very little overlap).

We then constructed spatial overlap networks visualizing the pairwise space-use overlap between all observed voles at each site using the ‘igraph’ R package (Csardi & Nepusz, 2006). Separate networks were constructed for each month June-October to capture the temporal dynamics of network structure and account for conspecific interactions that vary through time due to changing population size, demographics, or seasonal behaviors (Pinter-Wollman et al., 2014). In the networks, voles were represented as nodes and only voles captured in a given month were included in the network. Connections between nodes (edges) were undirected and were weighted by the value of pairwise space-use overlap between two voles (the edge weight between voles 1 and 2, £_1,2_) with no thresholding of edge weights.

#### 2.3.3 Individual spatial overlap

We quantified individual spatial overlap within the vole populations using weighted degree. Biologically, weighted degree represents the sum of all pairwise space-use overlap between a focal vole and its neighbors (“individual spatial overlap”). Values of weighted degree were unfiltered, allowing for very small measures of weighted degree indicating a very low probability of overlap (e.g., weighted degree <0.01). Two additional measures of spatial overlap: unweighted degree and normalized unweighted degree, are presented in the Supplement: Individual spatial overlap - Additional metrics (Table S1; Fig. S1, Fig. S2). All network metrics were calculated using the ‘igraph’ R package (Csardi & Nepusz, 2006).

Network data are inherently non-independent and degree is influenced by network size (total number of nodes in the network), limiting the ability to make direct comparisons between networks of varying size (Farine & Whitehead, 2015). Network size varied between sites and across months based on the number of voles observed during a trapping occasion. For an analysis of the effects of treatment on network size, see Supplement: Network size (Table S2).

## 3 RESULTS

We captured voles and recorded sex and reproductive status data for 742 unique individuals during our study period (June-October 2021). Of these, 445 voles (60%) were captured at least twice (mean ± SD captures per vole: 2.75 ± 2.33 [range: 1-17 captures]). We documented 905 captures across the 12 sites in summer (June-August) and 1107 captures in autumn (September-October). These capture data were used to inform our seasonal space use models.

Fecal egg count data was collected for 1035 captures during the study period. Of these, helminth eggs were detected in 440 samples (42.5%). Treatment group (control/deworm) had a significant effect on the relationship between pre-and post-treatment levels of helminth prevalence (odds ratio=0.51, p=0.022; Table S3) and infection intensity (/3=-0.74, p=0.005; Table S4). Helminth prevalence and the infection intensity were similar pre-treatment and both were lower in treated voles compared to control voles post-treatment. In dewormed voles, helminth prevalence remained similar from pre-treatment to post-treatment whereas helminth prevalence increased in control voles (Fig. S3A). Helminth intensity decreased in dewormed voles post-treatment while intensity was similar pre-and post-treatment in control voles (Fig. S3B).

### 3.1 Seasonal space use

In summer, the seasonal space use model indicated that the effect of distance on capture probability differed by vole sex, reproductive status, and site-level food supplementation treatment (GLM interaction terms between each parameter and distance from centroid: all p<0.001; Table S5; Fig. S4A-C). No effect of helminth removal was detected (GLM interaction term between helminth treatment and distance from centroid: p=0.18, Table S5; Fig. S4D).

When we estimated summer space use (using Eqn. 1) for each functional group in each treatment, we found evidence for male voles having greater space use than females and reproductive voles having greater space use than non-reproductive voles (Fig. 2). Food supplementation generally decreased vole space use, compared to unfed treatments (Fig. 2).

**Figure 2.**
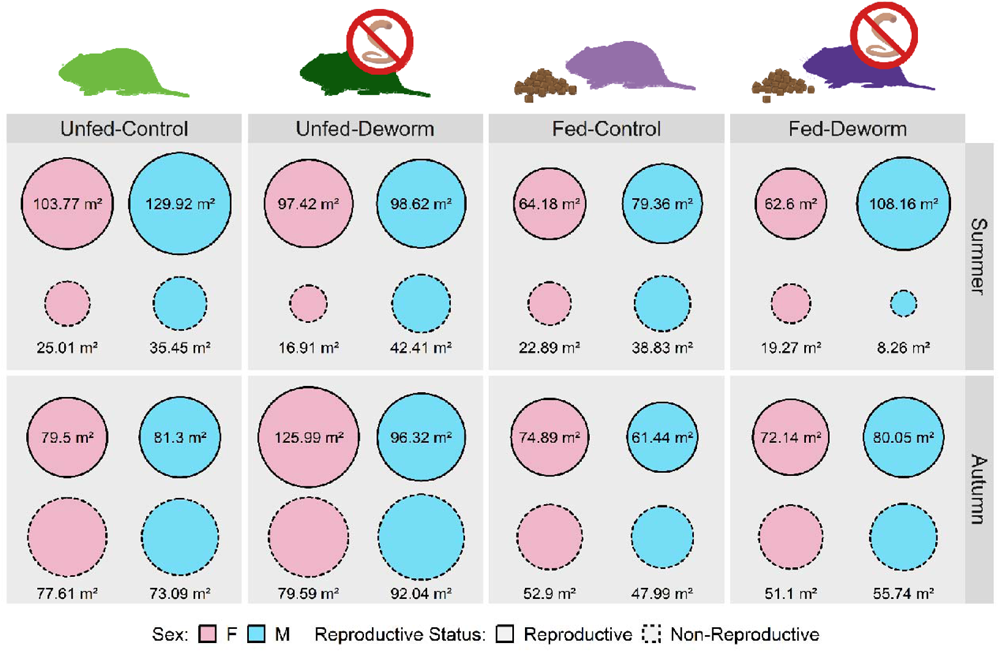
Space use of bank voles by sex and reproductive status across treatments in the summer breeding season (June-August) and autumn nonbreeding season (September-October). Circle size represents the mean space use area (m^2^) where the probability of capture is >0.01.

In autumn, the seasonal space use model indicated that the effect of distance on capture probability differed by vole reproductive status, site-level food treatment, and helminth treatment (GLM interaction terms between each parameter and distance from centroid: all p<0.012, Table S6; Fig. S5B-D). No effect of vole sex was detected in autumn (GLM interaction term between sex and distance from centroid p=0.68, Table S6; Fig. S5A).

When estimating space use, we found that reproductive voles had greater space use than non-reproductive voles, though space use of reproductive and non-reproductive voles was more similar in autumn than in summer (Fig. 2). The space use of reproductive voles generally decreased or remained similar from summer to autumn. The space use of non-reproductive voles increased from summer to autumn. Food supplementation decreased vole space use compared to unfed treatments (Fig. 2). Helminth removal had the opposite effect: voles in deworm treatments had increased space use, compared to voles in control treatments (Fig. 2).

### 3.2. Spatial overlap networks

In every month, all voles overlapped with at least one other vole and populations appeared densely and homogeneously connected if all edges were included in the network (i.e., no edge weight thresholding). However, edges of low weight (<0.05) were common and more heterogeneous network structure appeared with minimal edge weight thresholding (Fig. 3). Network size was highly variable across months and treatments (range: 2-52 voles). Networks were smallest in June, peaked in size in August (unfed treatments) or September (fed treatments), and then decreased slightly though October (Fig. 4A; see also Supplement: Network Size).

**Figure 3.**
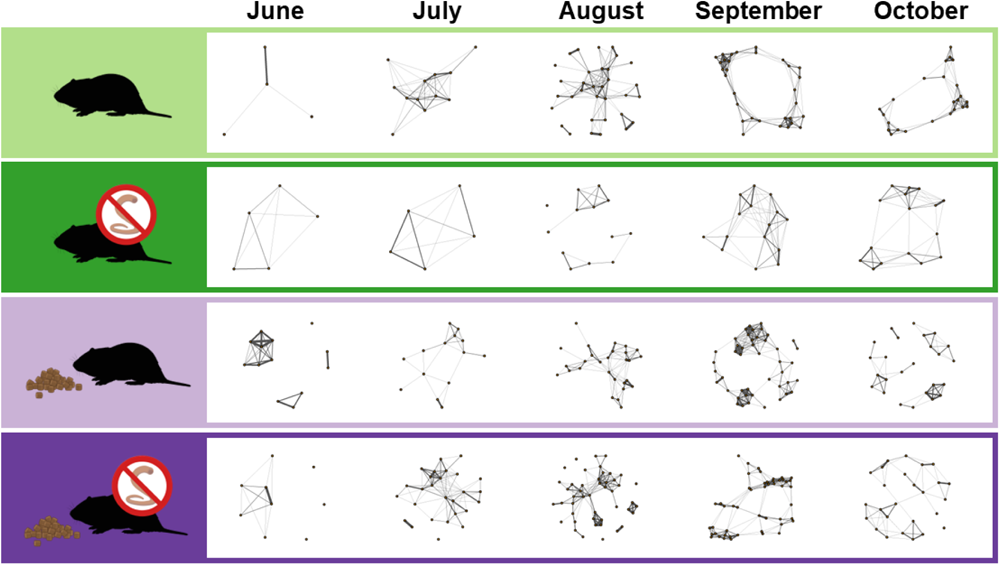
Spatial overlap networks of vole pairwise space-use overlap. Nodes represent individual voles, edges represent pairwise space-use overlap, thicker edges indicate greater space-use overlap between voles. For each treatment, networks from one site are shown. Edge weights are thresholded to a minimum edge weight of 0.05 for visualization purposes.

**Figure 4.**
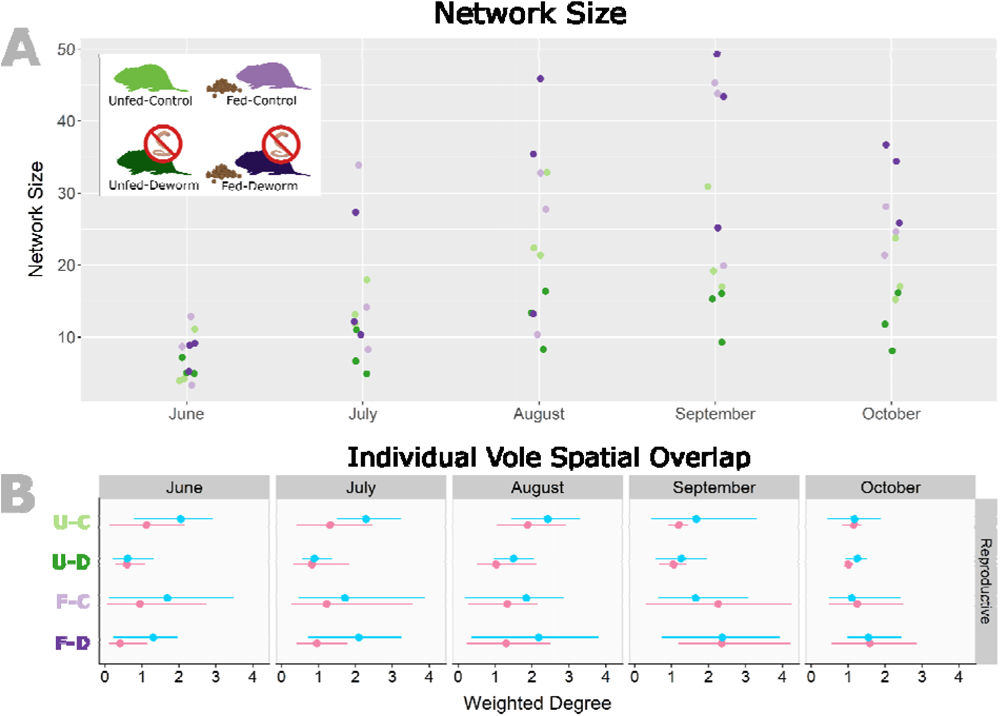
Population-level network size and individual vole spatial overlap. (A) Network size (number of voles) by month. One data point shown per replicate site. (B) Weighted degree (total pairwise space-use overlap per vole, weighted by the amount of overlap with each of its neighbors) by month. Means shown as points and lines indicate the range of values for males (blue) and females (pink) in each treatment. Data shown for reproductive voles only, trends for non-reproductive voles were similar.

Mean weighted degree, indicating individual vole spatial overlap (total spatial overlap of a focal vole with its neighbors, weighted by the amount of pairwise space-use overlap with each neighbor) was similar between the unfed-control, fed-control, and fed-deworm treatments in a given month in both summer and autumn. The range of weighted degree values were widest in September when the most voles with high degree (weighted degree >3) were observed (Fig. 4B). Compared to the other treatments, mean weighted degree in the unfed-deworm treatment was smaller with a narrower range in June-September (Fig. 4B). In summer (June-August), mean weighted degree was generally greater for reproductive male voles than reproductive females. In autumn (September-October), weighted degree was more similar between reproductive male and female voles (Fig. 4B). Similar trends were observed for non-reproductive voles.

## 4 DISCUSSION

Variation in food availability and macroparasite infection can alter animal space use and intraspecific interactions in ways that influence spatial processes in wildlife. Through experimental food supplementation and helminth removal in wild bank vole populations, we quantified the effects of environmental and individual variation on space use and spatial overlap. Food supplementation decreased vole space use, while helminth removal increased vole space use, but only in autumn. Space use was also influenced by functional group (combination of sex and reproductive status) and season. Mean individual vole spatial overlap was similar between treatments in a given month, despite up to three-fold differences in population density. Our findings show that both environmental variability and individual variation influence animal space use, but that space use and population density can trade off and maintain consistent spatial overlap in wildlife populations under different environmental conditions.

We detected significant effects of both food supplementation and deworming treatment on vole seasonal space use, though the effect sizes were generally small. The experimental and replicated design of our study and the large seasonal sample size of captured animals likely enables the detection of subtle and potentially important differences in space use. Food supplementation decreased vole space use in both summer and autumn. These findings align with the general understanding that supplemental food decreases bank vole home range size (Bondrup-Nielsen & Karlsson, 1985; Mazurkiewicz, 1983). However, food supplementation also increased population density, introducing a confounding factor which cannot be fully disentangled from the observed effects on space use. Increased population density can increase territorial behavior, resulting in smaller, more easily defended home ranges - potentially independent of food availability (Davis et al., 2015). Decreased space use under food supplementation is likely a complex response driven by interacting effects of food availability and population density.

In autumn, deworming treatment increased vole space use as compared to voles with unmanipulated helminth infection. This effect was not observed in summer, suggesting a time-lagged effect of deworming treatment. This may be due to individual-level factors; it takes time for the medication to start working and for behavioral effects of helminth infection to be impacted, so treatment in one month would not be observed to impact behavior until the following month. Moreover, the deworming treatment was only applied to captured voles and thus was heterogeneously applied across the population. As such, several months of trapping may have been necessary to treat a sufficient proportion of individuals and observe population-level effects (Pedersen & Fenton, 2015) or repeated treatments per individual were necessary to decrease the environmental burden and limit infection (Knowles et al., 2013). Increased space use by voles in the deworm treatments lends support to the hypothesis that animals with high helminth burdens exhibit sickness behavior and decrease their movements (Ghai et al., 2015). One potential confounding factor is that population density was very low across the unfed-deworm treatment, thus the increased space use may be a result of minimal territorial constraints and not a direct result of helminth removal. However, population sizes were more similar between the fed treatments (and, in some cases, populations were larger in fed-deworm than fed-control sites) and space use was still greater with deworming treatment, supporting helminth removal being a biologically meaningful factor influencing vole space use.

Space use interacted with vole sex and reproductive status as expected based on vole biology. We found that male space use was greater than that of females during the breeding season and space use of reproductive voles was greater than that of non-reproductive voles, which is consistent with the established literature (Bondrup-Nielsen & Karlsson, 1985; Bujalska, 1990). However, while we are confident in our space use estimates for reproductive voles, the space use of non-reproductive voles may be more difficult to estimate, particularly in summer. On average in the summer months, non-reproductive voles were captured less frequently and in fewer traps than reproductive voles (whereas capture frequency and trap usage were similar in autumn). We also observed very small space use by non-reproductive voles in summer. This could be representative of their true space use (e.g., a ‘sit and wait’ strategy, Bujalska, 1990), but may also be the result of many young voles who are captured once and then disperse off the trapping grid and are not captured again, artificially ‘shrinking’ the observed space use of non-reproductive voles. While we can not disentangle the relative impacts of these two factors, our observations still provide valuable comparisons of reproductive vole space use across seasons and environmental and individual variation by food supplementation and helminth removal.

It was surprising to us that, despite the differences in space use and population density, we did not observe clear differences in individual spatial overlap by treatment, even when population sizes differed. We found that food supplementation increased population density, consistent with previous manipulations in wild vole populations (Bujalska & Janion, 1981). It has been hypothesized that territorial behavior among reproductive females should be weakest when food is abundant and evenly distributed and population density is high (Ostfeld, 1985). Previous studies in *Clethrionomys* voles support this hypothesis, demonstrating that food addition decreases reproductive female space use, but increases spatial overlap between females (Andrzejewski & Mazurkiewicz, 1976; Ims, 1987). We observed decreases in both male and female space use with food supplementation, but did not observe an increase in spatial overlap in fed treatments compared to unfed. Similar spatial overlap despite differences in population density could be explained by a non-linear interaction of space use with population density. Individuals may move more freely and have larger space use at lower densities (unfed treatments), but constrain their space use and only contact their nearest neighbors at high densities (fed treatments; e.g., Davis et al., 2015). Thus, it is possible the effects of food supplementation on space use and population density may trade off, resulting in similar patterns of spatial overlap between fed and unfed populations, despite differences in population density.

Our network construction methods were chosen to minimize the effect of infrequent or missing observations of individuals on network structure. Networks in this study were constructed from 48 hours of observation with a possible maximum of four observations per vole and thus are not a complete representation of monthly space use. Networks constructed from trapping data often assign edges between individuals caught in the same or adjacent traps (e.g., Grear et al., 2009; Perkins et al., 2009), which can bias network measures if capture frequency is heterogeneous. We combined capture data across individuals and across months to define an average seasonal space use for each functional group. In this way, the behavior of animals captured many times fills in the ‘missing’ data for animals captured only once or twice and minimizes the effect of low capture frequency on network structure (Wanelik & Farine, 2022). Further, by using overlapping space-use to define connections in the spatial overlap networks, spatial overlap between voles is also less dependent on their specific trapped locations. These generalized approaches can help to minimize the uncertainty associated with network measures when social behavior is inferred using snapshot observations of individual behavior. One drawback is that aggregating behavior across functional groups eliminates individual variation within the group. Our objective was to quantify and compare space use between treatments and populations. By grouping voles by functional group, we allow behavior to vary across biologically meaningful axes such as sex and reproductive status (Henttonen, 2022), while capturing the average behavior of each population subclass which can be more readily generalized to other animal systems.

Spatial overlap can be linked to many direct and indirect interactions between animals including social interactions and pathogen transmission. When direct observation of interactions between animals is limited or impossible, thoughtfully aligned measures of spatial overlap can be used to approximate these interactions and predict transmission opportunities (Craft & Caillaud, 2011; Mistrick et al., 2022). However, spatial overlap and transmission opportunities may not correlate across all pathogen types. For instance, direct transmission can be decoupled from spatial overlap if animals seek out or avoid interactions with conspecifics. Thus, spatial overlap may be best used to approximate indirect transmission opportunities via the environment where direct contact between animals is not required (Godfrey et al., 2010). In that context, our finding that spatial overlap was similar across treatments could suggest that opportunities for environmental transmission may remain relatively constant even when population size increases–such as under high resource availability. However, differences in space use by sex and reproductive status could bias functional groups such as mature males toward greater exposure to environmental pathogens or greater potential to shed infectious material over larger areas (Krasnov et al., 2012). As such, the utilities of space use and spatial overlap data can be extended to various fields, providing insight into animal behavior and its ecological consequences, even when it cannot be measured directly.

## 5 CONCLUSIONS

Field studies investigating the effects of environmental and individual variation on animal spatial patterns are limited, thus this study provides necessary, empirical evidence of the effects of food supplementation and helminth removal on space use and spatial overlap in wildlife populations. We demonstrate that environmental variability due to supplemental food and individual variation in macroparasite infection help shape rodent space use. We also identify the potential for space use and population size to trade off, maintaining similar levels of spatial overlap between individuals despite variation in their environment. As such, our research demonstrates that environmental variability and individual variation drive animal spatial behavior and interact with population dynamics in wildlife populations and therefore represent important drivers of population-level contact-and proximity-driven processes such as social interactions and pathogen transmission.

## ACKNOWLEDGEMENTS

This research was funded by the National Science Foundation (DEB-1911925). J.M. was supported by the National Science Foundation Graduate Research Fellowship Program (award no. 2237827). Any opinions, findings, and conclusions or recommendations expressed in this material are those of the authors and do not necessarily reflect the views of the National Science Foundation. J.M. was also supported by the Dayton Fund of the Bell Museum and the University of Minnesota Ecology, Evolution, and Behavior Graduate Program and the College of Biological Sciences.

We are grateful to our collaborators at the Lammi Biological Station and University of Helsinki: John Loehr, Janne Sundell, Esa-Pekka Tuominen, Tiina Tulonen, Matti Kotakorpi, Riitta Ilola, Jaakko Vainionpää, Tomas Strandin, and Tarja Sironen, and to the students who helped collect and process the field data: Alexis Beagle, Muriel Chaudhri, Stephanie Du, Lucie Fornilli, Mathilde Gaudillère, Teemu Lemola, Nathaniel Mull, and Anni Simonen. Thanks to Vole Fever project collaborators Clay Cressler, Brendan Haile, and Lexi Beagle as well as Sharon Jansa and Susan Jones for providing feedback on this work.

Figures were created with BioRender.com. Fig. 1 site map by Anthony Shing.

## AUTHOR CONTRIBUTIONS

Kristian Forbes, Sarah Budischak, Meggan Craft, and Richard Hall conceived the ideas, designed the methodology, and secured the funding; Shannon Kitchen, Janine Mistrick, Jasmine Veitch, Samuel Clague, Brent Newman, Kristian Forbes, and Sarah Budischak collected the data; Janine Mistrick analyzed the data, created visualizations, and led the writing of the manuscript under the mentorship of Meggan Craft. All authors contributed critically to the drafts and gave final approval for publication.

## DATA AVAILABILITY STATEMENT

Data will be made available through the Dryad Digital Repository upon manuscript acceptance. Code is available in a GitHub repository (github.com/jmistrick/vole-spatial-v2) which will be archived on Zenodo upon manuscript acceptance.

