## Supplemental materials for "Effects of food supplementation and helminth removal on space use and spatial overlap in wild rodent populations"

SUPPORTING INFORMATION

**Individual spatial overlap - Additional metrics**

*Methods*

We explored individual spatial overlap within the vole populations using measures of weighted degree, unweighted degree, and normalized unweighted degree, calculated for each vole (Table S1).

**Table S1.** Description of network metrics and their biological relevance

| **Network Metric** | | **Network Definition** | **Biological Relevance** |
| --- | --- | --- | --- |
| **Node Level** | Weighted degree | Number of edges connected to a focal node by the weight of each edge; weights represent amount of spatial overlap | Sum of all pairwise space-use overlap a focal vole shared with its neighbors (“individual spatial overlap”) |
|  | Unweighted degree | Number of edges connected to a focal node | Number of unique neighbor voles with whom a focal vole overlapped |
|  | Normalized unweighted degree | Number of unweighted edges connected to a focal node divided by the number of nodes in the network less one; for comparison between networks of different sizes | Proportion of the voles in the observed population with whom a focal vole overlapped |
| **Network-Level** | Network size | Total number of nodes in the network | Number of voles observed during a trapping occasion at a given site |
|  | Degree distribution | Frequency distribution of degree counts in a network (weighted or unweighted) | Quantifies the heterogeneity in individual spatial overlap in the population |

Weighted degree (presented in the main text) was chosen to quantify the amount of overlap a focal vole had with its neighbors as a measure of cumulative opportunities for direct interactions or indirect exposure to a shared environment.

However, weighted degree does not capture the number of unique individuals a focal vole overlaps with, which could be important for transmission. We therefore also calculated unweighted degree (Table S1). For unweighted degree, thresholds of weighted degree (below which voles were considered not to meaningfully overlap) of 0.05, 0.01, 0.005, and 0.001 were investigated and the least restrictive threshold that resulted in voles with unweighted degree values of 0 was used to calculate unweighted degree (Fig. S1).

Population density was highly variable between treatments which would increase the maximum possible unweighted degree for an individual in a large population compared to a small population. To better compare the number of unique overlaps across populations of varying size, we also calculated unweighted degree, normalized by the population size. This measure therefore represents the proportion of the population with whom a focal vole overlapped in space.

Network data are inherently non-independent and degree is influenced by network size, limiting the ability to make direct comparisons between networks of varying size (Farine & Whitehead, 2015).The degree measures of all observed voles per treatment and month were visualized as degree distributions using density plots. In these, the x-axis indicates the value of interest, and the height of the curve suggests the frequency of occurrence of that value in the population. Narrow distributions suggest low variation in the population values while wide distributions indicate greater variation.

*Results*

Thresholds of weighted degree (weighted degree >0.05, >0.01, >0.005, >0.001) were investigated to choose an appropriate minimum value to constitute an overlap for unweighted degree. The threshold at 0.01 was chosen as a moderately restrictive threshold that resulted in some of the weakest overlaps having an unweighted degree of zero while still maintaining heterogeneity in the distribution of unweighted degree values (Fig. S1).

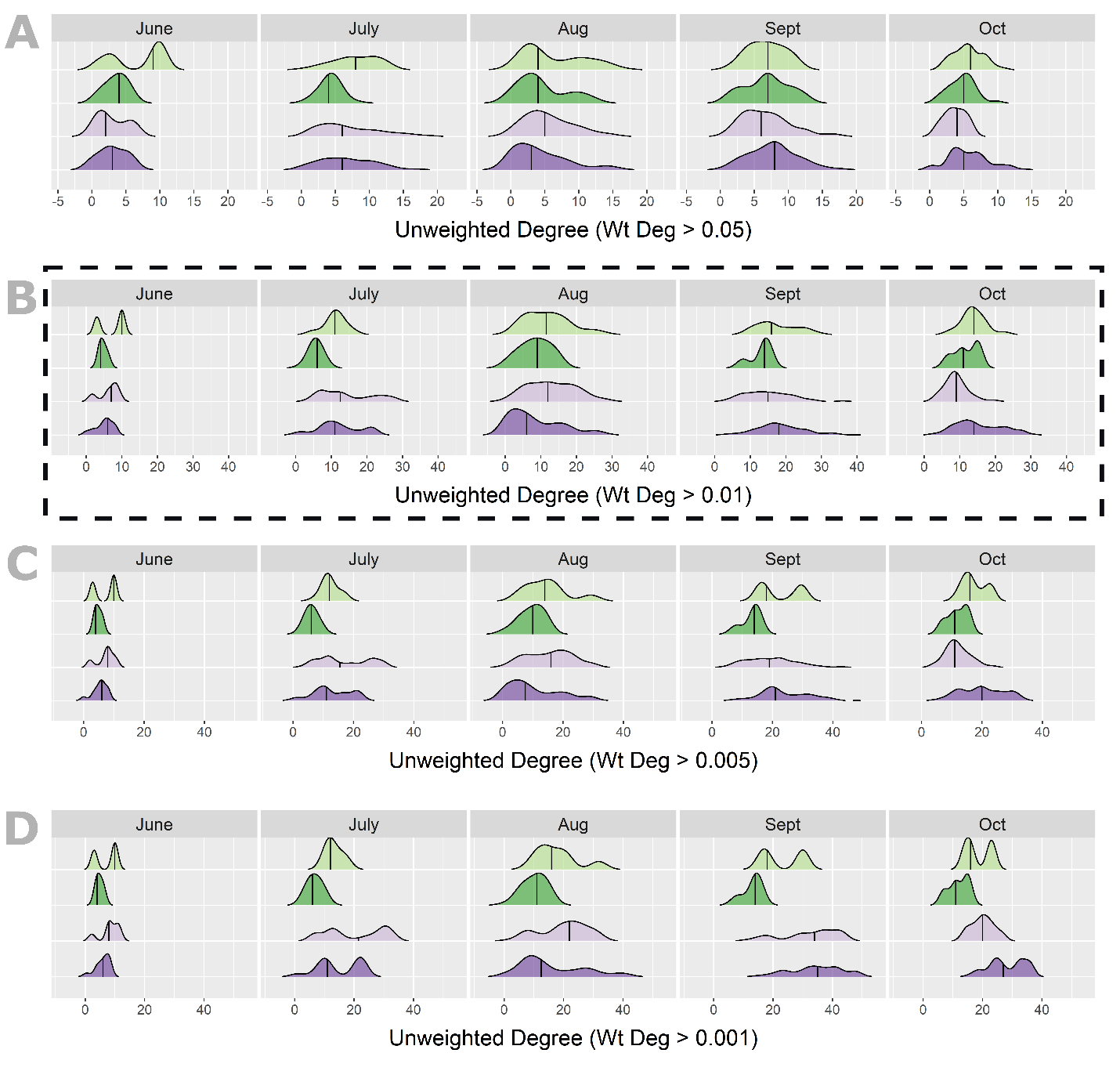

**Figure S1.** Sensitivity analysis of unweighted degree. Density distributions of unweighted degree with varying thresholds of weighted degree to constitute an overlap (A) weighted degree >0.05, (B) weighted degree >0.01, (C) weighted degree >0.005, and (D) weighted degree >0.001. Dotted line indicates the threshold used for analysis in the main text (weighted degree >0.01).

Distributions of unweighted degree (the number of unique neighbor voles with whom a focal vole overlapped, using a threshold of >0.01 to define an overlap) were generally wide and fairly similar between the unfed-control, fed-control, and fed-deworm treatments in a given month in summer and autumn (Fig. S2C). Unweighted degree in the unfed-deworm tended to have a more narrow distribution in most months. Like weighted degree, mean unweighted degree was similar across all four treatments in a given month.

Unweighted degree normalized by network size (i.e., the proportion of the population with whom a focal vole overlapped) was similar across treatments in June-August and showed wide distributions where voles overlapped with anywhere from 0-100% of the observed population. However, normalized unweighted degree was most different between treatments in autumn (Fig. S2D). In September and October, on average, voles in the unfed treatments overlapped with a majority to nearly all of observed population at their site (U-C: mean normalized unweighted degree 0.765 ± standard deviation 0.154; U-D: 0.990 ± 0.032) while voles in the fed treatments overlapped with less than half of the observed voles at their site (FC 0.381 ± 0.140; FD 0.470 ± 0.176).

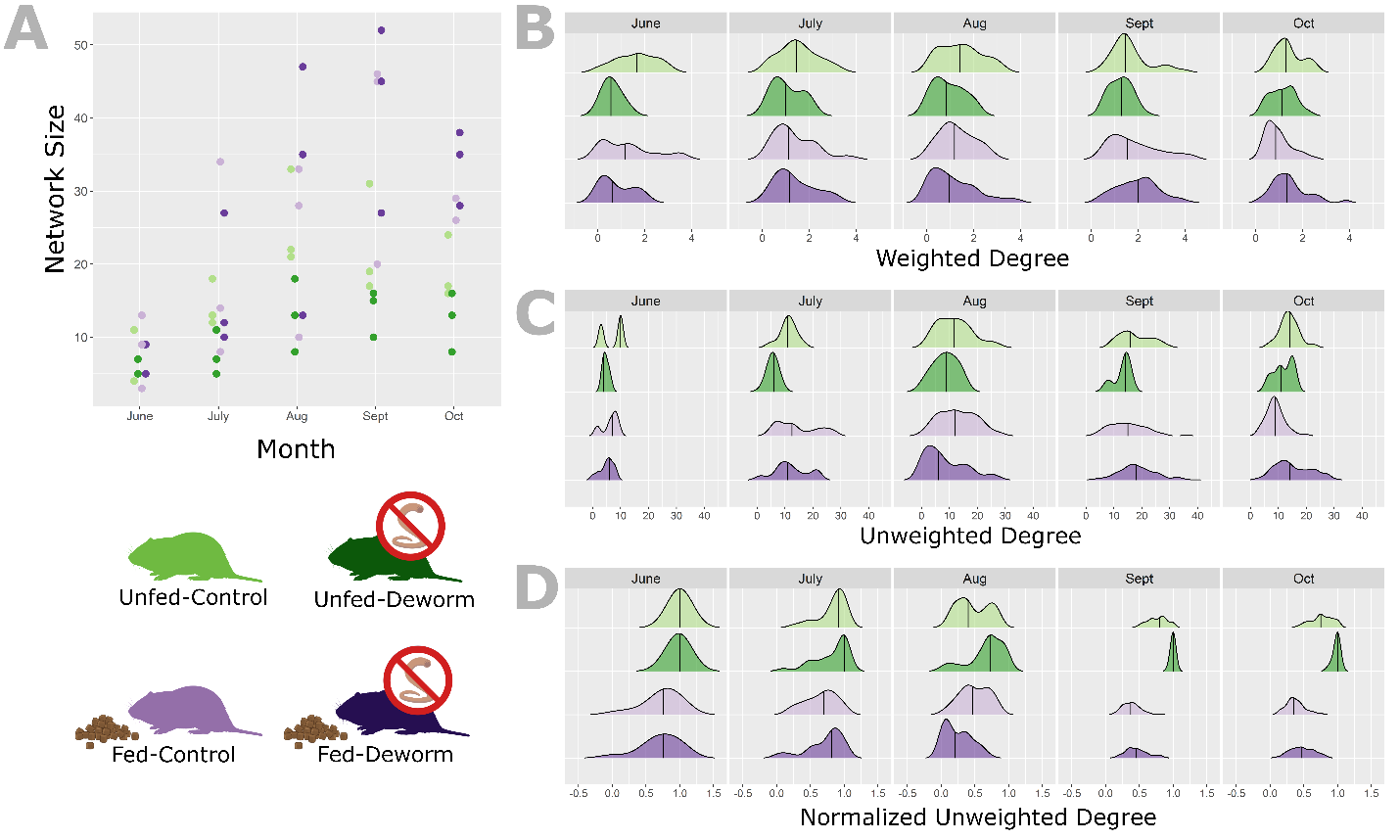

**Figure S2.** Network size and distributions of individual vole spatial overlap. (A) Network size (number of voles) by month. One data point shown per replicate study site (as shown in main text). Density distributions by month and treatment of three metrics quantifying individual spatial overlap (B) weighted degree (spatial overlap per vole, weighted by the amount of overlap with each of its neighbors; as shown in main text), (C) unweighted degree (number of unique neighbors each focal vole overlapped with where weighted degree ≥0.01), and (D) unweighted degree normalized by network size.

*Discussion*

The number of neighbor voles a focal vole overlapped with was highly variable among individuals, even within a treatment. The number of neighbors overlapped with per vole was most heterogeneous in fed treatments in September, which was also the month of highest population density. In September, individual voles in the fed treatments overlapped with anywhere from 0 to 40 other voles. There was less variability in the number of overlaps in September in the unfed treatments and all voles overlapped with at least three neighbors. High resource abundance clearly increased vole population density, increasing the potential maximum number of overlaps, but abundant resources may also enable voles to maintain sufficiently small territories and limit their overlap to only a few other animals (Ims, 1987). Particularly for functional groups such as reproductive females which exhibit territorial behavior, there may be increased pressure to maintain a strict territory when population density is high in order to protect offspring and maintain sufficient resources (Jonsson et al., 2002; Koskela et al., 1997). However, studies have also suggested that females may behave more amicably toward each other when resources are abundant, which would result in more overlap (Andrzejewski & Mazurkiewicz, 1976; Ims, 1987). Heterogeneity in individual spatial overlap could also be a product of foraging area, in conjunction or independent of territorial behavior, and voles may simply be able to forage in smaller areas when resources are abundant, limiting their potential for overlap with conspecifics.

The patterns observed in our study only suggest that the number of conspecifics overlapped with may be more heterogeneous in larger, food-supplemented populations. Nonetheless, this possibility does have implications for processes driven by spatial overlap such as pathogen transmission. Heterogeneous contact rates (either direct or via the environment) can generate superspreading events where a small number of individuals that are highly connected can drive transmission for the entire population (Lloyd-Smith et al., 2005). We may expect that high density populations such as those associated with food supplementation could be most likely to exhibit superspreading events. However, this would only be the case if a highly connected individual is infected. If an individual that overlaps with few other neighbors is infected, the pathogen would not spread as widely. As such, the possibility for sustained transmission and high population-level prevalence in large vole populations in autumn would be highly dependent on which individuals in the population were infected entering the autumn season.

The proportion of the population individual voles overlapped with was different between treatments in autumn. In unfed treatments in September and October, voles on average overlapped with 75-99% of the observed population, compared to 38-47% of the population in fed treatments where populations were much larger. This would further suggest that voles change their space use to maintain a similar absolute amount of spatial overlap, and not a similar proportion of neighbors overlapped with, when population density increases. While contact rates in populations are assumed to be either density-dependent or frequency dependent, population size and interactions between animals may not always scale, particularly if there are social behaviors limiting contacts (Cross et al., 2009). Our findings align with those of previous research in vole systems which indicate that contact-based transmission rates are a saturating function of host density (Smith et al., 2009), potentially indicating that contact rates are not as dichotomous as they seem. This is an exciting area of research which could benefit from future experimental studies to empirically test relationships between host density and contact patterns.

**Network size**

*Methods*

To understand how network size was affected by treatment and to contextualize potential confounding factors influencing degree measures, we investigated the factors affecting network size using a linear mixed-effects model fit by restricted maximum likelihood (REML) using the ‘lme4’ package (Bates et al., 2015). We modeled the main effects of food supplementation (unfed/fed), helminth removal (control/deworm), and month. Site was included as a random effect to account for multiple measures at each site and variation between replicate sites within a treatment. Network size was natural log-transformed to normalize the distribution of the data.

*Results*

Network size was moderately affected by food treatment (𝛽=0.48, *p*=0.022) and by month (𝛽=0.30, p<0.001; Table S2) with fed treatments and months later in the year having larger networks. Deworming treatment had no effect on network size (𝛽=-0.17, *p*=0.40; Table S2).
**Table S2.** Summary of linear mixed-effects model predicting natural log-transformed values of network size by food supplementation, helminth removal, and month. Significant p-values are bolded. Site ID is included as a random effect in all models (n = 12 sites, total observations = 60).

|  | **ln(Network Size)** | | | |
| --- | --- | --- | --- | --- |
| *Predictors* | *Estimate* | *Std. Error* | *CI* | *p* |
| (Intercept) | 1.96 | 0.19 | 1.58-2.35 | **<0.001** |
| Food Supplementation | 0.48 | 0.20 | 0.07-0.89 | **0.022** |
| Helminth Removal | -0.17 | 0.20 | -0.59-0.24 | 0.398 |
| Month | 0.30 | 0.04 | 0.22-0.38 | **<0.001** |
| **Random Effects** (Site ID) |  |  |  |  |
| σ^2^ | 0.18 |  |  |  |
| 𝜏_00 Site_ | 0.09 |  |  |  |
| Intra-class Correlation Coefficient _Site_ | 0.34 |  |  |  |
| N _Site_ | 12 |  |  |  |
| Observations | 60 |  |  |  |
| Marginal R^2^ / Conditional R^2^ | 0.482 / 0.659 | |  |  |

**Efficacy of deworm treatment**

**Table S3.** Summary of generalized linear mixed-effects model (binomial family, logit link) predicting the likelihood of helminth infection by treatment group, treatment stage, and their interaction. Vole ID is included as a random effect (n=701 individuals, n=1035 captures).

|  | **Helminth Infection** | | | |
| --- | --- | --- | --- | --- |
| *Predictors* | *Odds Ratio* | *CI* | | *p* |
| (Intercept) | 0.76 | 0.61 – 0.95 | | **0.016** |
| Treatment Group [Deworm] | 0.80 | 0.58 – 1.10 | | 0.173 |
| Treatment Stage [Post-treatment] | 1.58 | 1.07 – 2.33 | | **0.022** |
| Treatment Group * Treatment Stage  [Deworm] [Post-treatment] | 0.51 | 0.29 – 0.91 | | **0.022** |
| **Random Effects** (Vole ID) | | |  |  |
| σ^2^ | 3.29 | | | |
| τ_00_ _tag_ | 0.31 | | | |
| Intra-class Correlation Coefficient _tag_ | 0.09 | | | |
| N _tag_ | 701 | | | |
| Observations | 1035 | | | |
| Marginal R^2^ / Conditional R^2^ | 0.021 / 0.106 | | | |

**Table S4.** Summary of a linear mixed-effects model predicting helminth infection intensity: natural log-transformed eggs per gram of feces (EPG), by treatment group, treatment stage, and their interaction. Vole ID is included as a random effect (n=345 individuals, n=440 captures).

|  | **Helminth Intensity**  **ln(EPG)** | | | |
| --- | --- | --- | --- | --- |
| *Predictors* | *Estimate* | *CI* | | *p* |
| (Intercept) | 3.80 | 3.60 – 4.00 | | **<0.001** |
| Treatment Group [Deworm] | 0.10 | -0.20 – 0.40 | | 0.510 |
| Treatment Stage [Post-treatment] | 0.01 | -0.31 – 0.34 | | 0.935 |
| Treatment Group * Treatment Stage  [Deworm] [Post-treatment] | -0.74 | -1.26 – -0.22 | | **0.005** |
| **Random Effects** (Vole ID) | | |  |  |
| σ^2^ | 1.50 | | | |
| τ_00_ _tag_ | 0.17 | | | |
| Intra-class Correlation Coefficient _tag_ | 0.10 | | | |
| N _tag_ | 345 | | | |
| Observations | 440 | | | |
| Marginal R^2^ / Conditional R^2^ | 0.029 / 0.127 | | | |

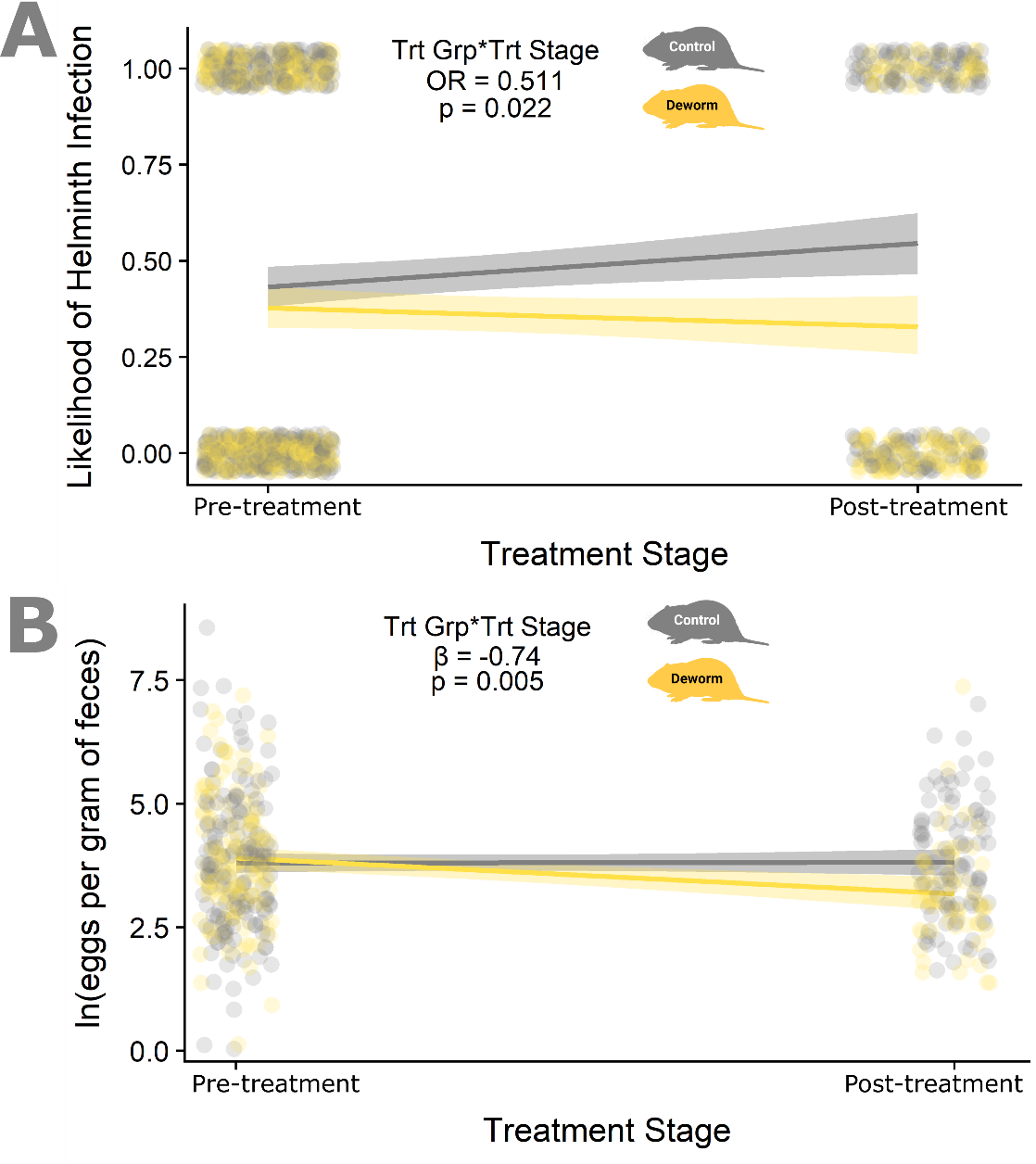

**Figure S3.** Visualization of mixed effects models quantifying the efficacy of the deworming treatment. Plots show the effects of treatment group (control/deworm) on the response variable from pre-treatment to post-treatment measurements. Modeled response variables are: A) helminth prevalence - using a generalized linear mixed-effects model to predict the likelihood of helminth infection and B) infection intensity - using a linear mixed-effects model to predict the natural log-transformed values of helminth eggs per gram of feces (EPG). Dots represent the raw data and lines indicate the fitted model for each treatment group. Model parameter estimates and p-values are reported for the interaction effect of treatment group*treatment stage in each model (see **Table S3:** helminth prevalence; **Table S4:** intensity).

**Space use**

**Table S5.** Model summary for the generalized linear model describing how the probability of capturing a vole varies with the natural logarithm of distance from its seasonal centroid, indicating vole space use, in the summer breeding season (June-August). Bolded p-values indicate significance.

| **Parameter** | **OR ^†^** | **95% CI ^‡^** | **p-value** |
| --- | --- | --- | --- |
| **Log Distance** | 0 | 0.00, 0.00 | **<0.001** |
| **Sex** |  |  |  |
| *Female* |  |  |  |
| *Male* | 0.21 | 0.12, 0.37 | **<0.001** |
| **Reproductive Status (seasonal)** |  |  |  |
| *Reproductive* |  |  |  |
| *Non-Reproductive* | 0 | 0.00, 0.00 | **<0.001** |
| **Food Treatment** |  |  |  |
| *Unfed* |  |  |  |
| *Fed* | 1.84 | 0.91, 3.80 | 0.093 |
| **Helminth Treatment** |  |  |  |
| *Control* |  |  |  |
| *Deworm* | 1.24 | 0.58, 2.68 | 0.58 |
| **Food Trt * Helm Trt** |  |  |  |
| *Fed * Deworm* | 0.57 | 0.20, 1.65 | 0.3 |
| **Log Dist. * Sex** |  |  |  |
| *Log Dist. * Male* | 4.59 | 2.70, 8.03 | **<0.001** |
| **Log Dist. * Reproductive Status** |  |  |  |
| *Log Dist. * Non-Reproductive* | 13,447 | 1,922, 121,475 | **<0.001** |
| **Log Dist. * Food Treatment** |  |  |  |
| *Log Dist. * Fed* | 0.31 | 0.16, 0.58 | **<0.001** |
| **Log Dist. * Helminth Treatment** |  |  |  |
| *Log Dist. * Deworm* | 0.65 | 0.33, 1.22 | 0.18 |
| **Log Dist. * Food Trt * Helm Trt** |  |  |  |
| *Log Dist. * Fed * Deworm* | 2.57 | 1.00, 6.74 | 0.053 |

^†^ OR - Odds Ratio	^‡^ CI - Confidence Interval

In summer, females were more likely to be captured within one trap of their centroid while males were more likely to be captured at greater distances (**Fig. S4A**). Non-reproductive voles were very likely to be captured only in one trap while reproductive voles were captured over a greater distance (**Fig. S4B**). Voles in unfed treatments were more likely to be captured 1-2 traps away from their centroid compared to voles in fed treatments (**Fig. S4C**). Voles in deworm and control helminth treatments were equally likely to be captured at all distances from their centroid (**Fig. S4D**)

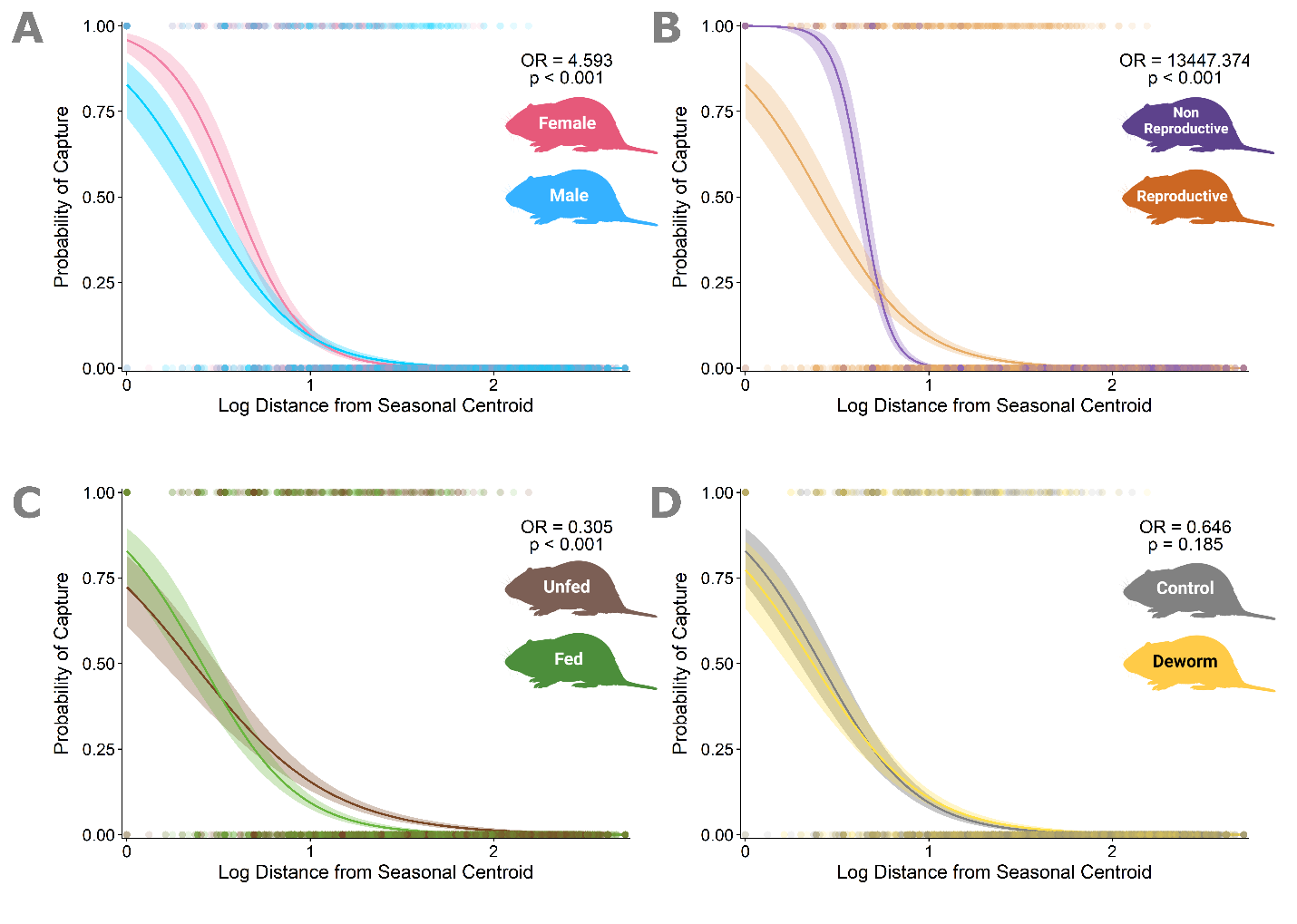

**Figure S4.** Subgroup-specific curves describing the change in capture probability with increasing distance from the seasonal centroid in the summer breeding season (June-August). Lines represent the fitted generalized linear model (GLM) for summer space use partitioned by each of the interaction terms in the model to show how the effect of distance on capture probability varies by (A) Sex: male vs. female bank voles, (B) Reproductive status: reproductive vs. non-reproductive bank voles, (C) Food treatment: voles in fed vs. unfed treatments, and (D) Helminth treatment: voles in deworm vs. control helminth infection treatments. Points show the raw data (whether a vole was captured at a given distance, 1, or not, 0). Distances are measured in trapping grid cells (1 grid cell=10 m). Odds ratios (OR) and p-values are reported for the interaction effect of each variable with the natural log of distance as modeled by the summer space use GLM (see **Table S5**).

**Table S6.** Model summary for the generalized linear model describing how probability of capturing a vole varies with the natural logarithm of distance from its seasonal centroid, indicating vole space use, in the autumn non-breeding season (September-October). Bolded p-values indicate significance.

| **Parameter** | **OR ^†^** | **95% CI ^‡^** | **p-value** |
| --- | --- | --- | --- |
| **Log Distance** | 0 | 0.00, 0.01 | **<0.001** |
| **Sex** |  |  |  |
| *Female* |  |  |  |
| *Male* | 0.75 | 0.42, 1.34 | 0.34 |
| **Reproductive Status (seasonal)** |  |  |  |
| *Reproductive* |  |  |  |
| *Non-Reproductive* | 0.61 | 0.33, 1.11 | 0.1 |
| **Food Treatment** |  |  |  |
| *Unfed* |  |  |  |
| *Fed* | 2.15 | 0.91, 5.11 | 0.080 |
| **Helminth Treatment** |  |  |  |
| *Control* |  |  |  |
| *Deworm* | 0.43 | 0.19, 0.97 | **0.043** |
| **Food Trt * Helm Trt** |  |  |  |
| *Fed * Deworm* | 1.91 | 0.58, 6.20 | 0.28 |
| **Log Dist. * Sex** |  |  |  |
| *Log Dist. * Male* | 1.13 | 0.64, 2.02 | 0.68 |
| **Log Dist. * Reproductive Status** |  |  |  |
| *Log Dist. * Reproductive* | 2.46 | 1.37, 4.38 | **0.002** |
| **Log Dist. * Food Treatment** |  |  |  |
| *Log Dist. * Fed* | 0.24 | 0.10, 0.57 | **0.001** |
| **Log Dist. * Helminth Treatment** |  |  |  |
| *Log Dist. * Deworm* | 2.61 | 1.23, 5.54 | **0.012** |
| **Log Dist. * Food Trt * Helm Trt** |  |  |  |
| *Log Dist. * Fed * Deworm* | 0.55 | 0.17, 1.81 | 0.33 |

^†^ OR - Odds Ratio	^‡^ CI - Confidence Interval
In autumn, males and females were equally likely to be captured at a given distance from their seasonal centroid (**Fig. S5A**). Non-reproductive and reproductive voles were equally likely to be captured within a trap distance from their centroid but reproductive voles were more likely to be captured further than one trap (**Fig. S5B**). Voles in the fed and unfed treatments were also equally likely to be captured less than one trap from their centroid, but voles in the unfed treatments were more likely than fed treatment voles to be captured at least one trap from their centroid (**Fig. S5C**). The difference was subtle between voles in the control and deworm treatments, but dewormed voles were more likely to be captured more than one trap from their seasonal centroid, compared to voles with unmanipulated helminth infections (**Fig. S5D**).

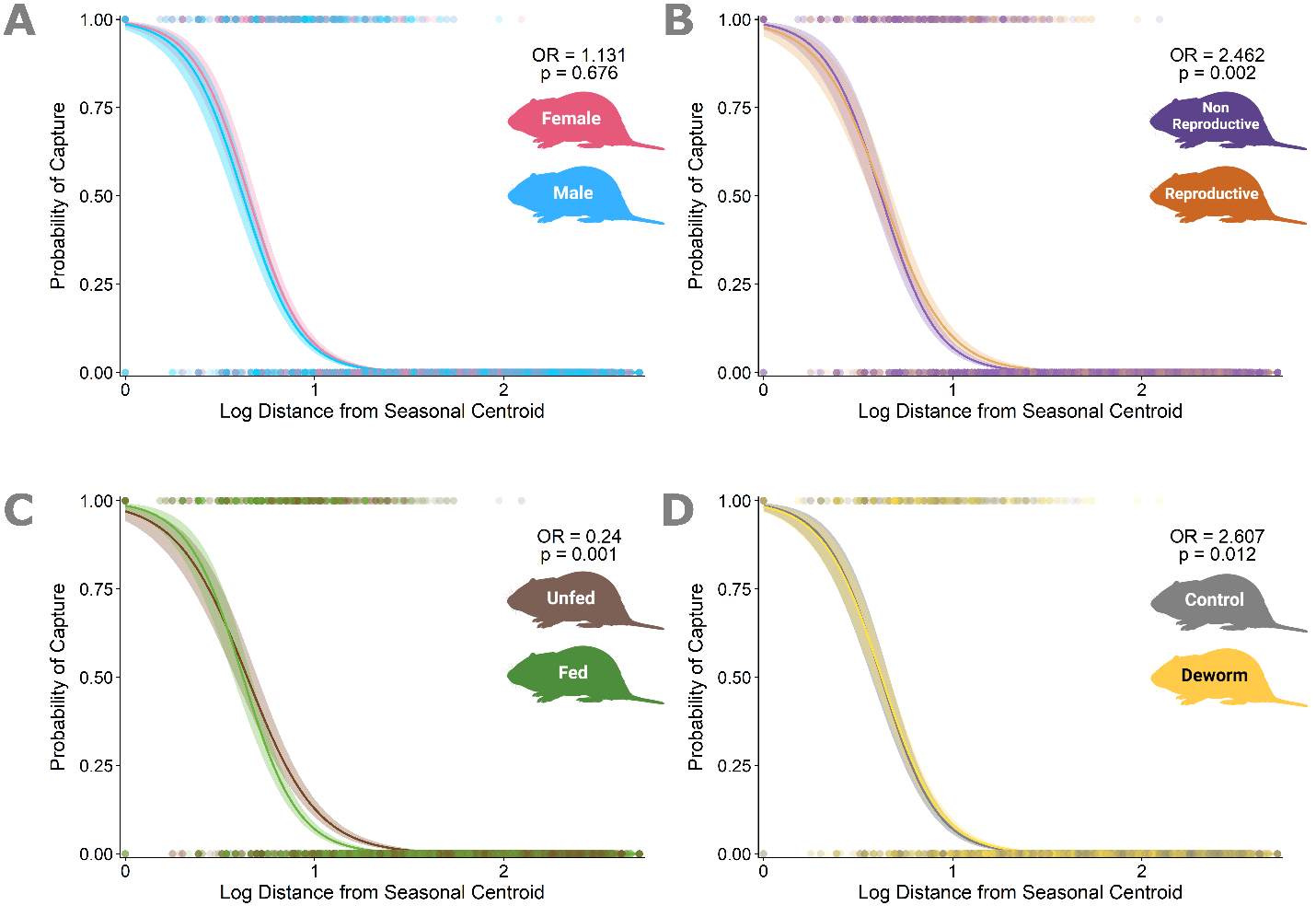

**Figure S5.** Subgroup-specific curves describing the change in capture probability with increasing distance from the seasonal centroid in the autumn non-breeding season (September-October). Lines represent the fitted generalized linear model (GLM) for autumn space use partitioned by each of the interaction terms in the model to show how the effect of distance on capture probability varies by (A) Sex: male vs. female bank voles, (B) Reproductive status: reproductive vs. non-reproductive bank voles, (C) Food treatment: voles in fed vs. unfed treatments, and D) Helminth treatment: voles in deworm vs. control helminth infection treatments. Points show the raw data (whether a vole was captured at a given distance, 1, or not, 0). Distances are measured in trapping grid cells (1 grid cell=10 m). Odds ratios (OR) and p-values are reported for the interaction effect of each variable with the natural log of distance as modeled by the autumn space use GLM (see **Table S6**).
